# Functional Interrogation of Lynch Syndrome Associated *MSH2* Missense Variants Using CRISPR-Cas9 Gene Editing in Human Embryonic Stem Cells

**DOI:** 10.1101/459586

**Authors:** Abhijit Rath, Akriti Mishra, Victoria Duque Ferreira, Chaoran Hu, James P. Grady, Christopher D. Heinen

**Author notes:** Corresponding Author: Christopher D. Heinen, 263 Farmington Avenue, Farmington, CT 06030-3101.

## Abstract

Lynch syndrome (LS) is a hereditary cancer predisposition condition caused by inactivating germline mutations in one of the DNA mismatch repair (MMR) genes. Identifying a deleterious germline mutation by DNA sequencing is important for confirming an LS diagnosis. Frameshift and nonsense mutations significantly alter the protein product and likely impair MMR function. However, the implication of a missense mutation is often difficult to interpret. Referred to as variants of uncertain significance (VUS), their discovery hampers the definitive LS diagnosis. To determine the pathogenic significance of a VUS it is helpful to know its impact on protein function. Functional studies in the test tube and in cellular models have been performed for some VUS, however, these studies have been limited by the artificial nature of the assays. We report here an improved functional assay in which we engineered site-specific *MSH2* VUS using Clustered Regularly Interspaced Short Palindromic Repeats (CRISPR)-Cas9 gene editing in human embryonic stem cells. This approach introduces the variant into the endogenous *MSH2* loci, while simultaneously eliminating the wild-type gene. We then characterized the impact of the variants on cellular MMR functions including DNA damage response signaling upon challenge with a DNA alkylating agent and the repair of DNA microsatellites. We classified the MMR functional capability of 8 of 10 VUS under study providing valuable information for determining their likelihood of being *bona fide* LS mutations. This improved human cell-based assay system for functionally testing MMR gene VUS will facilitate the identification of high risk LS patients.

**Significance Statement:** Understanding how cancer-associated missense variants in MMR genes affect function helps determine whether they truly contribute to disease. Laboratory assays previously utilized are limited by their artificial nature. To improve this, we introduced variants directly into the endogenous MMR loci in hESCs using CRISPR-Cas9 gene editing. This approach allows us to assess each variant while being expressed by its normal regulatory elements in a cellular environment. Our results will help guide the management of patients world-wide who carry these variants. At the same time, this study provides a technical road map for assessing the functional effects of all LS-associated variants, as well as variants linked to other genetic diseases where a cell-based functional assay is available.

## Introduction

DNA mismatch repair (MMR) performs multiple important cellular functions to protect genomic integrity. The primary function is to repair DNA polymerase errors. Recognition of the misincorporated base is carried out by the heterodimers composed of Mutator S homologs (MSH) MSH2-MSH6 or MSH2-MSH3. Upon detection of a mismatch, the MSH complexes recruit the Mutator L homolog (MLH) heterodimer MLH1-PMS2 which together regulate the activity of the DNA exonuclease Exo1 to excise the newly synthesized DNA strand prior to DNA re-synthesis (1, 2). MMR also attempts to repair mismatches introduced by polymerase mispairing of modified bases due to exposure to certain DNA damaging agents such as DNA alkylating agents. Futile repair processing of these lesions ultimately results in cell cycle arrest and apoptosis which prevents subsequent accumulation of mutation (3–5). In the absence of functional MMR, cells have an increased rate of spontaneous mutation generation and heightened resistance to the cytotoxic effects mediated by certain DNA damaging agents (6). Mutations in the MMR genes are associated with Lynch syndrome (LS), a hereditary cancer predisposition condition. LS is inherited in an autosomal dominant fashion with patients inheriting a mutation in one allele of a MMR gene. Somatic loss of the functioning wild type (WT) allele leads to a MMR deficient cell and the establishment of a well-characterized mutator phenotype. The increased mutation rate increases the likelihood of acquiring mutations in tumor suppressors or oncogenes ultimately leading to development of cancer (7).

Among the four canonical MMR genes, germline mutations in *MSH2* account for 35-40% of LS cases (8). Apart from the clearly deleterious changes such as nonsense or frameshift mutations that result in the loss of the MSH2 protein, a significant portion (~ 32%) of LS-associated *MSH2* variants are missense variants that result in the change of a single amino acid (9). For a large subset of missense variants, it has been difficult to unambiguously ascertain their impact on MSH2 function and therefore their significance to disease pathogenesis. Thus, they are aptly termed variants of uncertain significance (VUS). To help clinicians definitively diagnose suspected LS patients carrying germline VUS, a group of scientists and clinicians have created a 5-point classification scheme for MMR variants (10). The classification scheme attempts to determine the extent of pathogenic significance of a given variant based on co-segregation with disease, occurrence in multiple LS families, molecular characteristics of the tumor, and other features. In addition, determining whether the variant affects the function of the predicted protein product in a laboratory assay is often a crucial piece of information in determining its pathogenic likeliness (10).

The impact of *MSH2* VUS on protein function has been examined by multiple assays (11). *In vitro* reconstitution of the MMR reaction with cellular extracts or recombinant proteins, ectopic expression of the variant protein in MMR-deficient cancer cells, or modeling mutations in conserved yeast or mice *Msh2* residues have been utilized (12–15). However, possible caveats in these studies such as lack of a cellular environment, non-physiological level of mutant protein expression, or species-specific differences reduce confidence in their outcome. To this end, we have employed CRISPR-Cas9 as a tool to model a panel of LS-associated *MSH2* VUS in human embryonic stem cells (hESCs). As a non-transformed cell system hESCs provide several advantages. Unlike commonly used cancer cell lines, hESCs are an immortalized yet genetically stable, isogenic population. We created a panel of cell lines each harboring a specific *MSH2* variant in homozygous fashion and tested their ability to perform MMR cellular functions including repair and damage response signaling. These proof-of-principle experiments establish hESCs as a novel and valid cellular model to study the functional significance of LS-associated VUS in order to improve their clinical interpretation and better identify at-risk LS patients.

## Results

### Generation of *MSH2* VUS Expressing Cell Lines

We selected ten *MSH2* VUS as characterized by the International Society for Gastrointestinal Hereditary Tumors (InSiGHT) database to determine their effects on cellular MMR function (10) (**Fig. 1A**). We also selected two variants previously determined to be benign polymorphisms and two others previously deemed pathogenic mutations as positive and negative controls, respectively. We also used the parental H1 hESCs and an H1 hESC clone in which we used CRISPR-Cas9 to disrupt the *MSH2* loci in a homozygous fashion as described previously (16) as additional positive and negative controls, respectively. To assess the functional effects of *MSH2* VUS on MMR function in a human cell culture model, we used CRISPR-Cas9 gene editing to introduce each variant directly into the endogenous *MSH2* locus of H1 hESCs. A plasmid vector-based expression of both guide RNA (gRNA) and *Streptococcus pyogenes* Cas9 (SpCas9) was used to introduce a genomic DNA double-strand break (DSB) at the appropriate location in *MSH2.* We then selected clones which successfully utilized homology directed repair (HDR) using a single strand deoxyoligonucleotide (ssODN) as the external template to site-specifically introduce the mutations in a homozygous fashion (**Fig. S1, Table S1**). To enhance the efficient generation of single-cell derived hESC clones carrying the desired mutation, we employed several strategies. First, silent mutations were incorporated in the ssODN template along with the target amino acid-altering mutations to prevent Cas9 from re-cleaving after HDR of the break site. In some cases, these silent mutations also provided us with a way to screen for the mutant clone using restriction enzyme (RE) digestion of the PCR amplicon (**Fig. S2**). Second, SCR-7 (DNA ligase IV inhibitor) and L-755507 (Rad51 activity stimulator) were used during the targeting process to reduce non-homologous end joining repair of the DSB and increase the HDR efficiency (17, 18). Third, ssODNs with canonical 5’-PO_4_ and 3’-OH ends were used for increasing the incorporation efficiency into genomic DNA (19). The results of our gene editing experiments are summarized in **Table 1** and **Fig. S3** which show successful generation of 14 different *MSH2* homozygous variant-expressing cell lines.

**Fig. 1.**
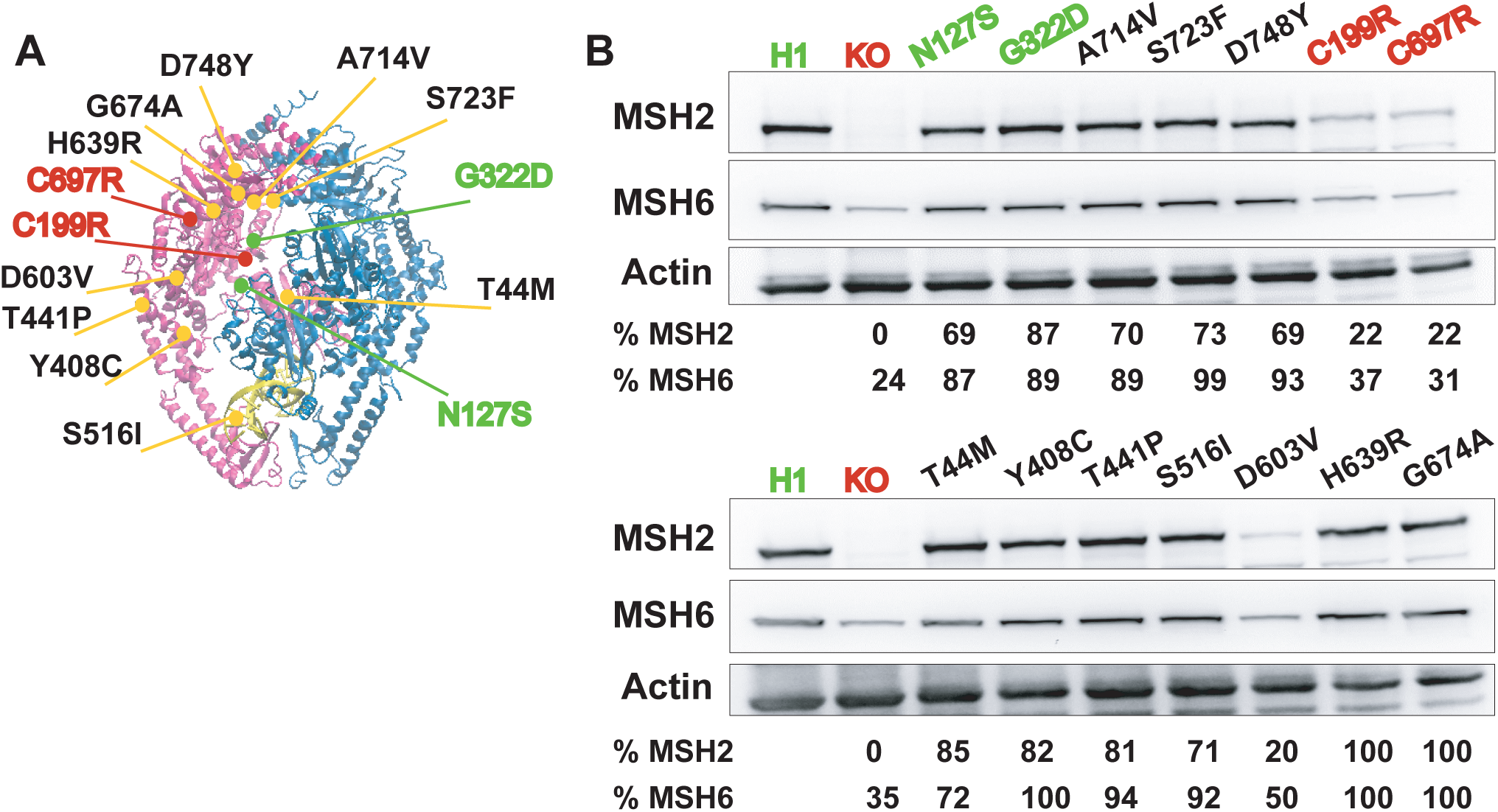
MSH2 variants tested and quantification of MSH2 and MSH6 steady-state protein levels. (*A*) The *MSH2* missense variants examined in this study as mapped to the DNA bound MSH2- mSh6 crystal structure (39). DNA is shown in yellow. (*B*) Steady-state level of MSH2 and MSH6 proteins in each of the engineered *MSH2* variant expressing lines is shown. Wild type and MSH2 Class 1 polymorphisms are depicted in green. *MSH2* KO and Class 5 pathogenic variants are shown in red. Actin was used as a loading control. Relative MSH2 and MSH6 levels were calculated with respect to the level in H1 WT cells after normalization for total protein loading.

**Table 1.** Targeting efficiency and relative position of target codon to PAM site

| Cell Line | No. of homozygous clones obtained | No. of clones screened | Homozygous targeting efficiency (%) | Distance of codon-altering mutations from PAM | Distance of silent mutations from PAM |
| --- | --- | --- | --- | --- | --- |
| *Class 1* |  |  |  |  |  |
| N127S | 2 | 24 | 8<sup>d</sup> | 4, 5 | 3, 6 |
| G322D | 1 | 7 | 14<sup>e</sup> | PAM | - |
| *Class 3* |  |  |  |  |  |
| T44M | 3 | 32 | 9<sup>d</sup> | 7 | 3, 9 |
| Y408C | 2 | 16 | 13<sup>e</sup> | 3 | - |
| T441P | 5 | 53 | 9<sup>d</sup> | 4 | 1,5 |
| S516I | 2 | 16 | 13<sup>e</sup> | 1 | - |
| D603V<sup>a</sup> | 0 | 45 | 0<sup>d</sup> | 31 | 3, 4, 5, 6, 21 |
| D603V<sup>b</sup> | 0<sup>c</sup> | 94 | 0<sup>e</sup> | 0 | - |
| H639R | 4 | 23 | 17<sup>d</sup> | 4 | PAM, 3, 5 |
| G674A | 1 | 22 | 5<sup>d</sup> | PAM | 4 (3' of PAM) |
| A714V | 3 | 11 | 27<sup>e</sup> | 1 | - |
| S723F | 2 | 35 | 6<sup>d</sup> | PAM | 2 |
| D748Y | 0<sup>c</sup> | 23 | 0<sup>d</sup> | 14 | PAM, 2, 6, 8 |
| *Class 5* |  |  |  |  |  |
| C199R | 1 | 16 | 6<sup>e</sup> | 7 | - |
| C697R | 5 | 48 | 10<sup>d</sup> | 1 | PAM, 3 |

<sup>a</sup> Initial attempt to target via an NGG PAM site failed to produce any targeted clones
<sup>b</sup> Targeting via an NGA PAM site
<sup>c</sup> A heterozygous clone was obtained that was re-targeted to generate the homozygous line
<sup>d</sup> Determined by restriction enzyme digestion screening
<sup>e</sup> Determined by Sanger sequencing

As shown in **Table 1**, we identified homozygous clones with efficiencies ranging from 5-27% as determined by restriction enzyme (RE) digestion screening, when applicable. To further confirm the incorporation of the target variant in a homozygous fashion, we employed Sanger sequencing. In certain instances, sequencing revealed that the positive result in the RE digestion screen only indicated presence of the silent mutation and not the target variant of interest. As an example, we identified two potentially targeted clones for the D748Y variant using the RE digestion assay. However, upon sequencing, we noted that the amino acid-altering mutation was only incorporated in a heterozygous fashion (D748Y_het_). Fortuitously, homozygous incorporation of the silent mutations provided a new PAM and gRNA binding sequence closer to the target codon. Thus, D748Y_het_ was further re-targeted using a re-designed gRNA that bound the modified protospacer region with the already incorporated silent mutations to generate a homozygous D748Y line (**Fig. S4**).

For a subset of variants where the PAM site was in close enough proximity to the target codon, we were able to introduce the target variant with relatively high efficiency without introducing additional silent mutations. If the target variant occurs in the PAM proximal regions (< 10 nt), then it should be sufficient to prevent re-cutting while simultaneously allowing for efficient targeting as Cas9 is intolerant to mismatches in this region (20–23). As shown in **Table 1**, we generated homozygous clones confirmed by Sanger sequencing at a modestly improved efficiency (compared to the lines in which additional silent mutations were included) for all such variants except C199R. Of note, the efficiency of single nucleotide insertion was higher for the variants within 1-3 nt of 5’ end of the PAM sequence (12-27%). However, an increase in distance, as in C199R (7 nt away from PAM), significantly decreased the targeting efficiency (6%).

Availability of a canonical PAM sequence for Cas9 in the genomic region of interest is a known limiting factor for a desired targeting event. In our study, we generally observed a relationship between the efficiency of targeting and the distance of the target codon from the PAM site, in agreement with previously reported observations (21, 24). Not surprisingly, the variants most difficult to target were the aforementioned D748Y which had a distance of 14 nt between target codon and its closest PAM site and the D603V variant which was 31 nt away from an available PAM. For D603V, we did not obtain a single homozygous clone initially as judged from the RE digestion assay (**Table 1**). Thus, we redesigned a gRNA using an NGA sequence which has been reported previously to be an alternate PAM site for SpCas9 (25, 26). We were able to successfully generate a single heterozygous clone which was subsequently re targeted to obtain multiple D603V homozygous clones (**Fig. S5**).

An important concern during CRISPR-Cas9 gene editing is off-target binding of the gRNA and the possibility of introducing a DSB by Cas9 mediated cleavage at unintended loci. To check for evidence of off-target cleavage in all 14 single-cell derived lines, we identified the top five predicted off-target sites by *in silico* analysis based on the probability of binding of a given gRNA to other locations in the genome. For each cell line, genomic DNA was extracted and the putative off-target regions were amplified by PCR. Purified PCR amplicons were subjected to Sanger sequencing and matched with the H1 hESC genomic reference sequence. We did not detect any sequence alterations at these sites in the genome suggesting an absence of off-target cleavage (**Table S2**) in line with similar observations from previous studies (27, 28). Taken together, these results demonstrate the practicality of using CRISPR-Cas9 as a versatile tool to engineer multiple, site-specific LS-associated *MSH2* VUS in an isogenic cell system to further analyze their functional impact on MMR function.

### Variant Effects on MSH2 Protein Level

Many pathogenic variants of *MSH2* impair MMR by conferring a defect in the stability of MSH2 protein (6, 11). To check the steady-state level of MSH2 in the different variant expressing lines, we measured MSH2 protein levels in comparison to the levels in WT H1 hESCs via Western blot. The known pathogenic variants C199R and C697R showed an approximately 80% reduction in MSH2 protein levels compared to the WT control. However, with the exception of D603V (20% of WT levels), we did not observe any appreciable changes in the MSH2 level of any other variant expressing lines (**Fig. 1B**). To determine if any of the variants interfered with the ability of the MSH2 protein to form a heterodimer with its partner MMR protein MSH6, we also examined MSH6 protein levels. MSH6 is an obligate heterodimer partner of MSH2 and so the inability of the two to form a heterodimer should lead to reduced stability of MSH6 (29). We found that the MSH6 protein levels were stable for all variant expressing lines except for those in which the MSH2 protein was also reduced suggesting that none of these variants likely interfere with heterodimer formation.

### DNA Damage Response Signaling

To test the functional effects of the altered MSH2 proteins in hESCs, we next examined the ability of these cells to elicit a DNA damage response to the alkylating agent MNNG. Human cancer cells undergo a permanent cell cycle arrest in response to various DNA damaging agents including S_N_1 alkylating agents like MNNG, however, this response is dependent on a functional MMR pathway (5, 6). More recently, we have demonstrated that human pluripotent stem cells instigate a rapid apoptotic response to MNNG, also in a MMR-dependent manner (16, 30). To assess the ability of the *MSH2* VUS lines to induce cell death in response to MNNG, we used an MTT assay to measure cell survival following treatment. A 24 h challenge with MNNG resulted in a significant survival advantage for *MSH2* knock-out hESCs compared to WT cells. Approximately 85% of the knock-out cells survived treatment at the highest concentration of MNNG tested (2 μM) compared to only 35% survival for WT cells (**Fig. 2A**). As expected, the response of hESC lines expressing known *MSH2* polymorphisms (N127S and G322D) and *MSH2* pathogenic mutations (C199R and C697R) matched closely to the WT and *MSH2* knock-out control cell lines, respectively (**Fig. 2A**). For the ten *MSH2* VUS expressing lines under study, we used a statistical clustering approach to identify those variants whose survival curves formed a separate subgroup from the pathogenic-like or benign-like controls. We created three, non-overlapping homogenous clusters with one based on data from the “pathogenic” standards, a second created from the “benign” standards, and a third intermediate cluster that was not similar to either of these groups. Using this statistical method, we were able to segregate the ten variant lines into the three clearly distinct phenotypic groups. T44M, Y408C, T441P, and A714V exhibited a similar response to that of the “benign” controls. The D603V, G674A, S723F, and D748Y cell lines were resistant to MNNG and clustered with the “pathogenic” controls. However, the S516I and H639R lines showed an intermediate phenotype upon MNNG challenge (66% and 67% survival, respectively at 2 μM MNNG) that formed an independent non-overlapping “intermediate” cluster (**Fig. 2A**). We reasoned that the impaired damage signaling in the S516I variant line should be rescued by reverting the mutant isoleucine residue back to serine. As we did not destroy the Cas9 target site during the initial targeting event, we were able to use CRISPR-Cas9 to correct the introduced single nucleotide variant and generate an I516S line. As shown in **Fig. 2B**, reversion of the mutant residue back to the original sequence restored MNNG induced apoptotic signaling to WT levels. Taken together, these results show the ability of our cell based assay system to clearly distinguish phenotypic variation between different *MSH2* VUS lines.

**Fig. 2.**
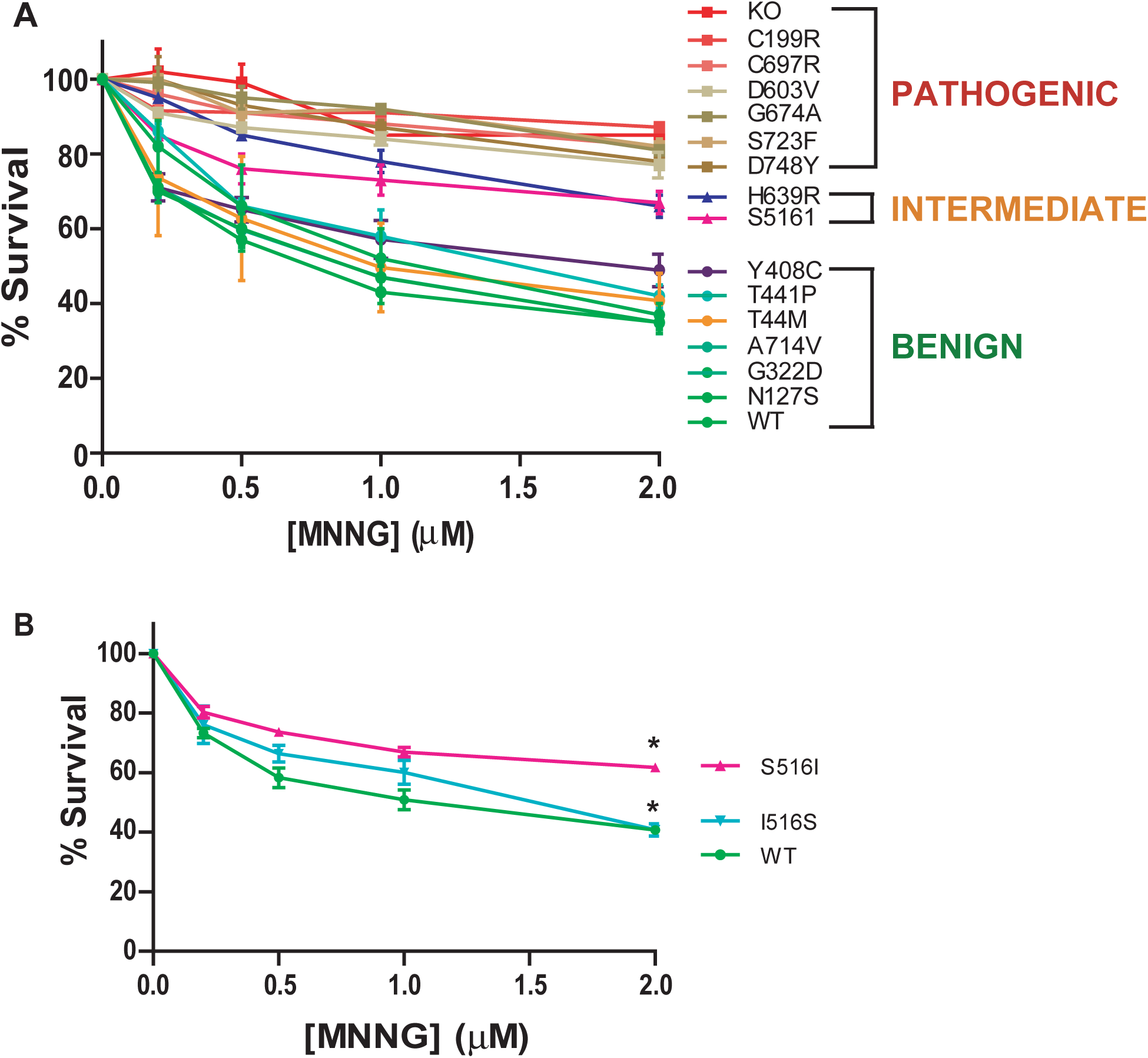
The MMR dependent damage response in variant cell lines. Cell survival as measured by MTT assay showing percentage survival after 24 h treatment with increasing doses of the DNA alkylating agent MNNG in the indicated cell lines. The values are represented as the mean ± S.E.M. N = 3-5. (A) Survival curves for 10 VUS lines along with WT and MSH2 knockout (KO) controls. A statistical clustering analysis was performed which grouped the variants into three populations based on their survival. One cluster resembled the “Pathogenic” controls, one cluster resembled the “Benign” controls and a third resembled an “Intermediate” cluster. (B) Survival curves for S516I line and I516S reversion mutant line. *, p < 0.05

### Mismatch Repair Ability

The canonical function of MMR is to detect and repair mismatches generated during DNA replication due to DNA polymerase errors (2). The MMR pathway also repairs small insertion and deletion loops created by polymerase errors at simple repeat sequences. To examine the ability of the MSH2 variants to repair endogenous chromosomal sequences, we tested for the presence of instability at two DNA microsatellites. Lack of functional MMR manifests as instability in genomic microsatellite regions which is the hallmark of *bona fide* LS tumors (31, 32). We tested for evidence of microsatellite instability (MSI) in the VUS expressing lines at two mononucleotide repeat loci, BAT-26 and NR-27, which are well-established predictive biomarkers for impaired MMR function (33). Single-cell derived clones were grown from control and VUS cell populations using dilution cloning. The MSI status at the BAT-26 and NR-27 loci was assessed employing fragment analysis for multiple clones for each cell line. **Table 2** shows the percentage of clones showing a variable allele length compared to the reference H1 genomic sequence suggesting an alteration in number of repeat units (indicative of MSI) for both loci. As expected, no MSI was detected in any of the subclones for H1 WT hESCs at either marker. In contrast, every clone obtained from *MSH2* KO cells exhibited an altered repeat length. Inter-clonal variability with regards to repeat length was also observed among the *MSH2* KO clones (data not shown). Among the missense variants, no MSI was detected for T44M, Y408C, T441P, and A714V along with the polymorphism controls (N127S and G322D) indicative of retained normal MSH2 repair function. However, a varying degree of MSI was detected for D603V, G674A, S723F, and D748Y along with the pathogenic controls (C199R and C697R) suggesting different degrees of functional impairment in these lines. Interestingly, the H639R clones also did not exhibit any defect in MSI length whereas S516I displayed a very low percentage of cells with MSI despite their apparent functional defects in the alkylation damage response assay.

**Table 2.** Microsatellite instability in variant cell lines

| **Cell Line** | **BAT-26<sup>a</sup>** | **NR-27<sup>a</sup>** | **Clones Tested** |
| --- | --- | --- | --- |
| H1 (WT) | 0 | 0 | 24 |
| MSH2-KO | 100 | 100 | 24 |
| *Class 1* |  |  |  |
| N127S | 0 | 0 | 27 |
| G322D | 0 | 0 | 32 |
| *Class 3* |  |  |  |
| T44M | 0 | 0 | 32 |
| Y408C | 0 | 0 | 32 |
| T441P | 0 | 0 | 32 |
| S516I | 19 | 6 | 32 |
| D603V | 71 | 60 | 31 |
| H639R | 0 | 0 | 32 |
| G674A | 52 | 26 | 31 |
| A714V | 0 | 0 | 32 |
| S723F | 41 | 25 | 32 |
| D748Y | 85 | 88 | 26 |
| *Class 5* |  |  |  |
| C199R | 34 | 53 | 32 |
| C697R | 100 | 100 | 20 |

<sup>a</sup> Percentage of clones with altered alleles compared to WT sequence

## Discussion

Limited genetic and functional information on a MMR gene missense variant can be a bottleneck in the definitive diagnosis of LS. In absence of a robust family pedigree, variant specific functional information becomes crucial in determining the likely pathogenic significance of the variant (34). Thus, a suitable model system to study the functional impact of a gene variant is necessary. The advent of CRISPR-Cas9 gene editing provides the ability to engineer specific genetic variants at their endogenous chromosomal locus. Thus, any impact of a given variant on protein function can be examined in the context of the gene’s cell-intrinsic regulatory mechanisms that govern its expression and function. This approach resolves concerns that have arisen from previous functional studies of MMR VUS (11). *In vitro* reconstitution of repair of a single base pair mismatch has been a valuable tool to identify variants that clearly disrupt biochemical activity, however, this approach likely does not reveal all variants that could interfere with MMR activity *in vivo.* Additionally, cell based assays relying on exogenous expression of variant-containing transgenes provides cellular context, but may be prone to non specific effects due to improper regulation of transgene expression. More so, the availability of an established variant expressing mutant line opens up possibilities for modeling other potentially interesting aspects of the disease. For example, putative epistatic or synthetic lethal interactions can now be studied in the variant background. In addition, although outside the scope of this study, our use of hESCs allows for future studies to examine the ramifications of the variant in specific cell types. Pluripotency confers the ability to differentiate these cells into any tissue type in the body and hence provides potential avenues to carry out functional testing in a three-dimensional model system and, if need be, do drug testing as a proxy for the tissue of interest. Through the course of the current study, we have developed a robust CRISPR-Cas9 targeting methodology for use in human pluripotent stem cells which are generally not very amenable to genetic manipulations using HDR (24, 35, 36).

A key cellular function of MMR that we assessed was the ability of MSH2 to instigate apoptosis in response to DNA damage. This function is an important gate-keeping mechanism that prevents cells from going forward with unrepaired DNA lesions (4-6). Absence of this protective mechanism due to MMR deficiency may be an important indicator of the transformative potential of a cell (37). Based on the response of the variant lines to MNNG, we were able to cluster eight *MSH2* VUS into two distinct groups indicating presence or absence of a functional DNA damage signaling pathway (**Fig. 2A**). Of those “benign-like” lines (T44M, Y408C, T441P, and A714V), the steady state levels of MSH2 protein are comparable to WT cells. In the “pathogenic-like” line D603V, the MSH2 protein is present at about 20% of the WT level which may partially underlie the defective damage response. In the other three “pathogenic-like” lines (G674A, S723F and D748Y), while the protein levels appear stable, the variants affect residues in the ATPase domain of MSH2 spanning amino acids 620-855, which is crucial to MSH2 function. Thus, the impairment of MSH2 dependent signaling may be due to defective adenosine nucleotide processing (38). The S516I variant displays an “intermediate” response in the MNNG toxicity assay while displaying no observable difference in the MSH2 protein level compared to WT. S516I is present in the clamp domain of the MSH2 protein very near residues K512 and R524 which make significant, non-specific contacts with the DNA backbone upon mismatch binding (39). The change of a polar amino acid (serine) to a hydrophobic residue (isoleucine) at position 516 may affect these non-specific interactions to partially impair mismatch binding affinity. As shown in **Fig. 2B,** we were able to rescue the sensitivity of this variant to WT levels upon correcting the mutation in the variant line to re express S516 indicating the specific contribution of the isoleucine substitution to the observed MMR response before correction. The H639R variant also displays an intermediate damage signaling function. This variant lies within the ATPase domain of MSH2 as well as within a domain between amino acids 601-671 that has previously been shown to be important for Exo1 binding in a peptide-binding assay (40). Further biochemical studies will be required to ascertain the specific mechanistic defect in these mutant proteins.

We also assessed the MSI status of all the generated lines in this study. No MSI was detected for the T44M, Y408C, T441P, and A714V lines, which when combined with their normal damage signaling functions suggests that these four *MSH2* VUS do not alter MSH2 function and are therefore, likely non-pathogenic variants. In contrast, D603V, G674A, S723F, and D748Y display both an MSI and damage signaling phenotype closely matching that of the *MSH2* knock out and known pathogenic variants providing strong functional evidence that these are *bona fide* disease causing mutations. Curiously, the S516I and H639R variants show either only a minor MSI phenotype or no defect at all. Given their intermediate effects in damage signaling, these variants may be weak disease alleles, though additional testing is clearly warranted to more accurately gauge their impact on disease.

Traditionally functional evaluation of any gene variant is carried out in a post hoc fashion which is a time consuming process. Hence, to provide quick guidance as to whether a new found variant is deleterious or neutral, a number of *in silico* variant prediction platforms have been developed (41, 42). These platforms mainly predict the effect of a gene variant on protein function based on evolutionary conservation and amino acid charge. However, often times different prediction algorithms provide conflicting information. For example, N127S is a well-established polymorphic variant of *MSH2* which is also confirmed in this study to be MMR-proficient (10, 43). N127S along with G322D are also represented in the ExAC database (44) as known polymorphisms prevalent in the normal population. However, multiple variant predictive platforms identify N127S as deleterious in nature (**Table 3**). Similar observations were also made for T44M and A714V which were likewise shown to be MMR proficient in our assays. These results demonstrate the inaccuracy that still exists with such *in silico* approaches, and highlights the need for a robust dataset of functionally characterized variants for training the *in silico* platforms.

**Table 3.** Summary of functional studies and prior *in silico* predictions

| Cell Line | Protein Stability | Damage Response Signaling | DNA Repair Function | Polyphen 2 Prediction | Mutation Assessor Prediction |
| --- | --- | --- | --- | --- | --- |
| *Class 1* |  |  |  |  |  |
| N127S | + | + | + | Possibly Damaging | High |
| G322D | + | + | + | Benign | Medium |
| *Class 3* |  |  |  |  |  |
| T44M | + | + | + | Probably Damaging | High |
| Y408C | + | + | + | Probably Damaging | High |
| T441P | + | + | + | Benign | Low |
| S516I | + | +/- | +/- | Benign | Low |
| D603V | - | - | - | Probably Damaging | High |
| H639R | + | +/- | + | Probably Damaging | High |
| G674A | + | - | - | Probably Damaging | High |
| A714V | + | + | + | Probably Damaging | High |
| S723F | + | - | - | Probably Damaging | High |
| D748Y | + | - | - | Probably Damaging | High |
| *Class 5* |  |  |  |  |  |
| C199R | - | - | - | Probably Damaging | Medium |
| C697R | - | - | - | Probably Damaging | High |

In summary, we demonstrate the utility of using CRISPR-Cas9 genome-engineered hESCs to obtain relevant functional information for better clinical interpretation of LS associated, patient-derived *MSH2* VUS. Based on the gathered functional data, we were able to assign the functional capability of 8 of the 10 VUS (**Table 3**). This functional characterization satisfies an important criterion in the InSiGHT variant classification scheme for determining pathogenic potential (10).

## Materials and Methods

### Cell Lines

hESCs (H1) were obtained from the WiCell Research Institute and cultured on growth factor reduced Matrigel coated plates in hESC media (Peprotech) and passaged either by microdissection or by using StemPro Accutase Cell Dissociation Reagent (ThermoFisher Scientific).

### Generation of *MSH2* Variant Expressing Cell Lines

For each variant, a suitable gRNA sequence upstream of a 3’ protospacer adjacent motif (PAM; NGG/NGA for SpCas9 used in this study) in close proximity to the site of mutation (less than 10 nt) was chosen using an *in silico* program (crispr.mit.edu (21) and CRISPOR (45)) (**Table S1**). DNA oligos containing the gRNA sequence were cloned into the Px459V2.0 vector (Addgene, plasmid# 62988). One million H1 hESCs were pre-treated with ROCK inhibitor (Selleckchem) for 2 h and then transfected using Amaxa Stem Cell Nucleofector Kit 2 (Lonza) following the manufacturer recommended protocol using an Amaxa Nucleofector II. Two μg of the gRNA and SpCas9 expressing plasmid were transfected along with 2 μL of 100 μM ssODN (90 nt with 5’ PO_4_^-^ modification) (IDT) carrying the mutation to be incorporated and silent mutations (if any) for screening purposes **(Table S1)**. To increase the rate of homology directed repair, DNA ligase IV inhibitor, SCR-7 (Selleckchem), and Rad51 activator, L755507 (Selleckchem), were added the day of transfection (at 1 μM and 5 μM final concentration, respectively) for 48 h. Cells were treated with 1 μg/mL Puromycin (Sigma) starting 24 h after transfection for a period of 48 h to select for transfected cells. CloneR (STEMCELL Technologies) was used to promote survival and growth of single cell clones post selection. Surviving colonies were picked with a fraction of the cells used for preparing genomic hotshot DNA for screening by PCR and the remaining cells transferred to a Matrigel coated 96- well or 24-well plate for further passaging. Details for off-target analysis and screening are described in the SI Materials and Methods. Details of Western blotting, cell survival, microsatellite instability and statistical analyses can be found in the SI Materials and Methods.

## Acknowledgments

We would like to acknowledge technical support provided by Ms. Qingfen Yang and Mr. Chris Stoddard in preparing the reagents and providing helpful suggestions. This work was funded by the National Institutes of Health grant CA115783 and the State of Connecticut Regenerative Medicine Fund grant 13SCB-UCHC-06.

